# Robust evidence for modest neoselachian diversity loss across the K/Pg: Response to Guinot et al

**DOI:** 10.64898/2026.04.08.716966

**Authors:** Amanda Gardiner, Gregor H. Mathes, Rebecca Cooper, Kristína Kocáková, Jaime A. Villafaña, Daniele Silvestro, Catalina Pimiento

**Affiliations:** Department of Paleontology, University of Zurich; Zurich, 8006, Switzerland; Department of Genetics, University of Cambridge; Cambridge, CB2 3EH, UK; Chair of Physical Geography, University of Passau, Passau, 94032, Germany; Natural History Museum, University of Oslo, Oslo, 0562 Norway; Centre for Planetary Habitability, Department of Geosciences, University of Oslo, Oslo, 0315, Norway; Departamento de Ecología, Facultad de Ciencias, Universidad Católica de la Santísima Concepción, Concepción 4090541, Chile; Department of Biosystems Science and Engineering, ETH Zurich, Basel, 4056, Switzerland; Department of Biological and Environmental Sciences and Gothenburg Global Biodiversity Centre, University of Gothenburg, Gothenburg, 41319, Sweden; Department of Biosciences, Swansea University; Swansea, SA2 8PP, United Kingdom

## Abstract

We reconstructed the neoselachian diversity over the past 145 million years using a new occurrence dataset and DeepDive^1-3^. We recovered a small decline through the K/Pg following a steady increase during the Cretaceous, and a prolonged, substantial decline towards the present following a mid-Eocene peak^2^. Guinot et al. argue that our conclusions are compromised by problems in the underlying data and by the way extinction magnitude across the K/Pg was quantified. They cast doubt particularly on the pattern across the K/Pg, which they consider to be at odds with all previous analyses. We audited the alleged data issues reported in Guinot et al. and found that they include a combination of differences in data-treatment decisions, valid corrections and cases where our verification did not support their proposed changes (see Supplementary Information). We applied their data treatment and ran sensitivity tests to evaluate how this approach affects our results, specifically around the K/Pg. None of our tests recovered a catastrophic diversity loss for neoselachians during this interval. As such, we demonstrate that our findings are robust and consistent across different data treatments.

---

Although our findings contrast with previous works by Guinot et al.^4,5^, we note that recent independent analyses have also recovered a moderate, rather than catastrophic, diversity decline across the K/Pg^6,7^. Such results were found using an independently compiled dataset, different methodology, and were based on genus-level occurrences, which are more taxonomically stable than species-level. The absence of a diversity collapse across the K/Pg is further corroborated by a reconstruction of neoselachian diversification rates applying PyRate^8^ to our dataset, which showed that although neoselachians experienced extinction rates greatly exceeding background levels, these were accompanied by elevated origination rates, suggesting rapid diversity turnover^9^. Thus, while interpretations differ among studies, our results are consistent with recent research.

Guinot et al. argue that estimating diversity trajectories is an inappropriate methodology to understand the evolution of biodiversity in deep time. While we demonstrably appreciate the value of modelling speciation and extinction dynamics^8^, we emphasize that the reconstruction of diversity trajectories has greatly advanced our understanding of the evolution of life on Earth through hundreds of studies published over many decades. These include some of the most influential research in the field of paleobiology, which have revealed the rise and fall of major clades in the history of life^10-13^. Methods to infer past biodiversity have evolved, from statistical and Bayesian approaches to deep learning, leading to updated reconstructions of diversity dynamics^3,14-16^. We agree with Guinot et al. that “a multitude of evolutionary histories can produce similar diversity trajectories”. This is in line with the recent finding that the same diversity trajectory can be explained by multiple dynamics of speciation and extinction rates, which therefore become non-identifiable parameters in some cases^17^. Yet, this does not invalidate research on past biodiversity trajectories as a key biological indicator of macroevolutionary dynamics.

Importantly, our goal was to reconstruct biodiversity trajectories through time. We contend that Guinot et al. attempt to move the goalposts by focussing on extinction rate and magnitude. Furthermore, when estimating extinction across the K/Pg, they use raw taxonomic counts without accounting for biases inherent to the fossil record. However, up to 50–60% of apparent diversity changes in shallow marine fossils can be attributed to sampling biases, and even methods traditionally used to mitigate them fail to fully account for spatial and temporal heterogeneities^18,19^. Indeed, using uncorrected range-through estimates, we originally found a K/Pg decline of 32% ± 2%^2^. When explicitly accounting for multiple biases using DeepDive, we recovered a diversity trajectory displaying a ~10% drop in diversity across the K/Pg^2^, providing an analytically robust insight into neoselachian evolutionary history.

## Embracing the fossil record through analytical evaluation

Our dataset (herein, FINS) was designed as a transparent synthesis of current knowledge of the neoselachian fossil record, with the goal of representing the published literature consistently and reproducibly. As such, each occurrence is traceable to its primary source facilitating verification^1^. Furthermore, each taxonomic name is harmonised based on the taxonomy from Shark-References ^20^. In line with recent calls to promote open, transparent, and continuously updated biodiversity datasets^21^, FINS includes detailed documentation to enable its reuse, and allows users to contribute additional occurrences and to implement alternative taxonomic approaches^1^. Our ambition is therefore for this dataset to evolve as new discoveries are made and taxonomic interpretations are refined.

Guinot et al. list a number of issues with our data. The issues regarding uncertain nomenclature (~7% of all our data) highlight that identifications including qualifiers such as “cf.” and “aff” do not constitute definitive identifications and could potentially inflate age or geographic ranges. However, such qualifiers do not necessarily imply that the material is identifiable only at higher taxonomic levels, nor that it represents a different taxon. In palaeontological practice, open nomenclature often reflects caution in response to preservation limits or conservative identification standards, even when the material is considered closely comparable to the nominal species. Similarly, the suggestion to exclude records lacking illustrations or taxonomic descriptions (i.e., “without evidence”, ~40% of our dataset) imposes a highly restrictive data treatment that would remove substantial published information. Occurrence datasets commonly used in large-scale paleobiology research (e.g., ^22,23^) compile published, traceable reports of taxon presence rather than taxonomic diagnoses, with the effect of low-confidence data being testable. Indeed, previous empirical work has shown that even substantial errors in large taxonomic compilations do not bias broad diversity patterns when error is not systematic, underscoring the need to evaluate rather than assume its effect *a priori*^24^. Finally, the issues regarding age and taxonomy include both valid indications for corrections (~70% of cases) and claims that were proven unfounded after verification (~30%). Most corrections involved minor updates to taxonomic names, synonymies, or classifications and had limited impact on the composition of the analysed dataset. A detailed response to each issue is provided in the Supplementary Text and Table S1.

We applied the corrections deemed to be valid (Table S1) to FINS (Version v2), applied our filtering protocol^2^ (Data S1), and ran three sensitivity tests: 1) removing all occurrences with uncertain nomenclature (8% of occurrences removed from original dataset at the species level); 2) training DeepDive to explicitly model the K/Pg extinction; and 3) applying the same approach as the first test, but also removing occurrences “without evidence” (36% of occurrences at the species level; Table S2).

Under our first test, we recovered a diversity change of −22.9 ± 1.1% (Table S2; Data S2). Although this contrasts with the −10% ± 9% reported originally, it lies close to the upper range of the original estimate. Fundamentally, the magnitude still corresponds to a moderate decline rather than a catastrophic loss (Figure 1). Under our second test, which explicitly models the K/Pg extinction, we recovered a decline of −29.7 ± 8.4% and under our third test, which is the most restrictive, the magnitude of the decline remains within a comparable range (−33.1 ± 3.9%). Together, our sensitivity tests indicate that even under Guinot et al.’s data approach, the inferred species richness loss across the K/Pg remains moderate and incompatible with a catastrophic diversity decline. These values also contrast with those obtained based on uncorrected range-through, which range between 42-48% (Figure 1; Table S2; Data S3) further highlighting the effects of not explicitly accounting for spatial and temporal heterogeneities^18,19^.

**Figure 1.**
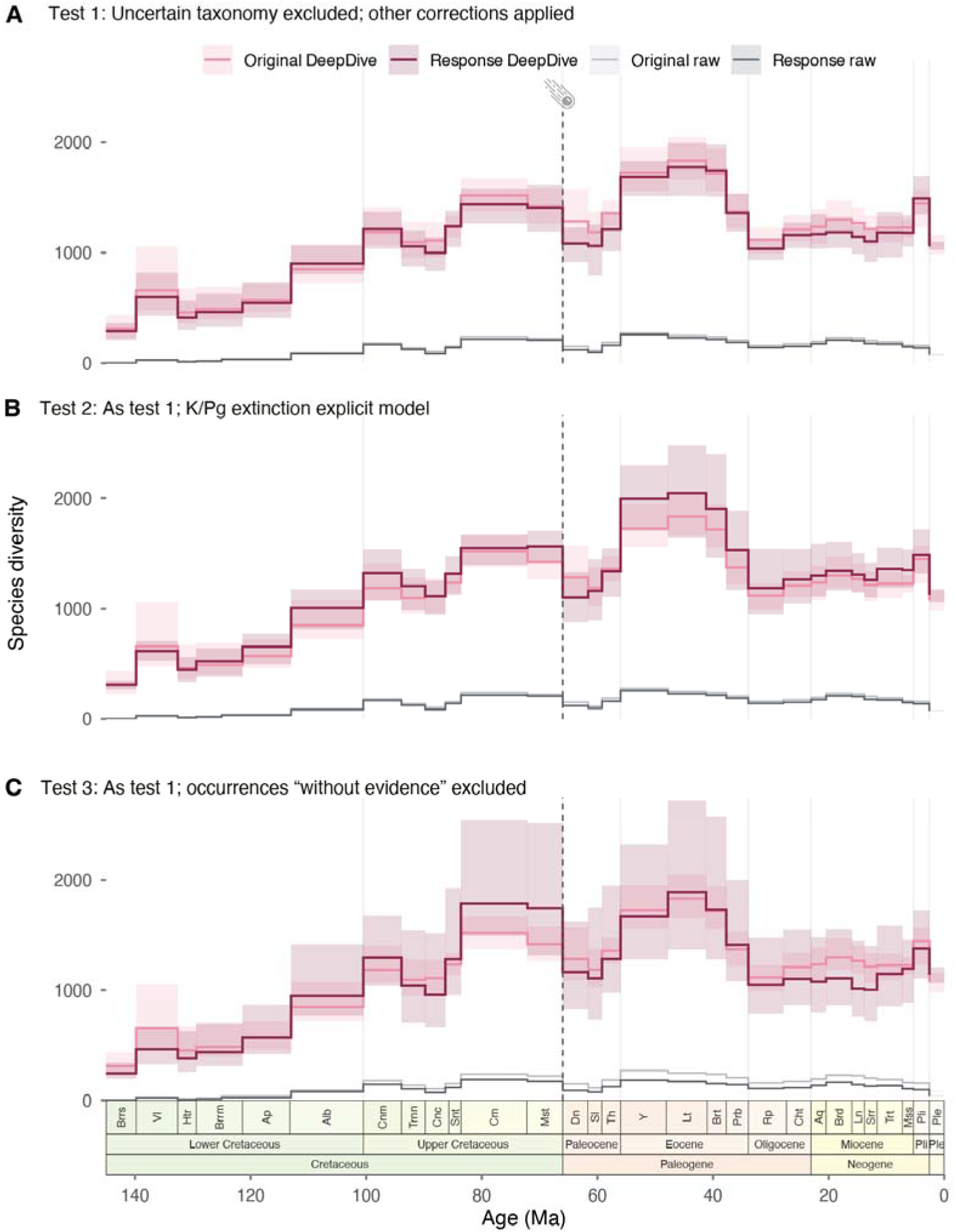
Species diversity trajectories reconstructed under alternative data treatments (Table S2). **A.** Sensitivity test 1, whereby we removed all occurrences with uncertain nomenclature, applied the corrections deemed to be valid after our own evaluation, and retained occurrences classified as “without evidence”. **B**. Same as A, but K/Pg extinction is explicitly modelled in the simulations. **C**. Same as A, but occurrences classified as “without evidence” are excluded. Solid lines represent the mean reconstructed richness across simulations, and shaded envelopes indicate the uncertainty around the estimates (maximum deviation between minimum and maximum predicted richness values across simulations). Vertical lines mark epoch boundaries. Abbreviations are as follows: Brrs, Berriasian; Vl, Valanginian; Htr, Hauterivian; Brrm, Barremian; Ap, Aptian; Alb, Albian; Cnm, Cenomanian; Trnn, Turonian; Cnc, Coniacian; Snt, Santonian; Cm, Campanian; Mst, Maastrichtian; Dn, Danian; Sl, Selandian; Th, Thanetian; Y, Ypresian; Lt, Lutetian; Brt, Bartonian; Prb, Priabonian; Rp, Rupelian; Cht, Chattian; Aq, Aquitanian; Brd, Burdigalian; Ln, Langhian; Srr, Serravallian; Trt, Tortonian; Mss, Messinian; Pli, Pliocene; Ple, Pleistocene. Dashed vertical line and meteorite shape denote the Cretaceous/Paleogene extinction event.

Although diversification rates were not the focus of our original paper, in response to Guinot et al.’s comment, we have applied the data treatments used in the sensitivity tests described above and estimated diversification trajectories using PyRate^8^. Across treatments, our findings align with previous diversification analyses^9^, showing that while extinction rates were high at the K/Pg (0.88-1.21), speciation rates also increased (0.31-0.87), yielding moderately negative net diversification estimates (-0.26 to -0.56; Fig. S1). These additional tests support our interpretation that the K/Pg signal reflects elevated turnover, with extinction buffered by origination, rather than a dramatic neoselachian diversity decline (Figure S1).

## Conclusions

Our evaluation of Guinot et al. revealed that the diversity trajectory originally recovered for neoselachians remains consistent (Figure 1), supporting a modest diversity decline through the K/Pg. The more restrictive data treatments explored here affect absolute estimates, but do not lead to inferring a catastrophic reduction in neoselachian diversity. These results align with independent diversity reconstructions using different approaches^6,7^, as well as diversification rate analyses suggesting rapid diversity turnover during this interval (Figure S1)^9^. Taken together, these multiple lines of evidence indicate that high origination rates buffered net loss through the K/Pg, resulting in a smaller diversity decline than previously thought^2,6,7,9^. Our results therefore do not deny elevated extinction across the K/Pg; rather, they add an important missing piece to the puzzle of neoselachian evolution.

## Resource availability

### Lead contact

Further information and requests for resources should be directed to and will be fulfilled by the lead contact, Catalina Pimiento.

### Materials availability

This study did not generate new unique reagents.

### Data and code availability

- The full FINS dataset used is accessible at Zenodo (Version v2): https://doi.org/10.5281/zenodo.20744849
- The filtered dataset used in the analyses is included in the Extended data (Data S1)
- All code and data used in this study are publicly available at Zenodo: https://doi.org/10.5281/zenodo.20735451. These include:

∘ The raw data
  ▪ The code to filter the FINS dataset
  ▪ The filtered dataset, divided in subsets for the analyses
  ▪ The paleo-oceans assigned to each occurrence and their time range

∘ The data and code to perform the DeepDive analyses:
  ▪ The codes to generate the input files to process the data in DeepDive
  ▪ Input files, including .csv occurrence data and .ini configuration files
  ▪ The codes to perform the DeepDive analyses
  ▪ The outputs from DeepDive

∘ The data and code to perform the uncorrected range-through (raw) analyses:
  ▪ The codes to perform the analyses
  ▪ The outputs

∘ The data and code to perform the PyRate diversification analyses:
  ▪ The codes to perform the analyses
  ▪ The outputs
▪ The codes and data to create all tables and figures included in this manuscript.

- The resulting diversity estimates over time using DeepDive at the species level are included in the Extended data (Data S2)
- The resulting diversity estimates over time using uncorrected range-through (raw) at the species level are included in the Extended data (Data S3)

## Acknowledgments

This work was supported by the Swiss National Science Foundation through a PRIMA grant (no. 185798) awarded to CP. DS received funding from ETH Zurich and the Swedish Foundation for Strategic Environmental Research MISTRA within the framework of the research programme BIOPATH (F 2022/1448).

## Supplemental Information

Supplemental information including procedures, figures, tables, and references can be found online.

