## Supplementary Information for "Robust evidence for modest neoselachian diversity loss across the K/Pg: Response to Guinot et al"

**Supplementary text**

FINS is an occurrence dataset that effectively synthesises the current knowledge on the neoselachian fossil record^1^. Each occurrence is linked to the primary literature, and each taxonomic name was harmonised based on the taxonomic dictionary from Shark References (<https://shark-references.com>, accessed in March 2021). In the version currently under review, we include a Shiny application that allows users to visualise and contribute additional occurrences and implement alternative taxonomic approaches. Our ambition is therefore for this dataset to evolve through time as new discoveries are made and taxonomic interpretations are refined. The dataset used for our analyses is a filtered version of FINS, which *only* includes: 1) occurrences identified to species- or genus-level; 2) taxonomic nomenclature and age deemed to be valid; and 3) occurrences with age ranges of less than or equal to 15 myr^2^.

***Assessment of the issues reported by Guinot et al.***

*Use of uncertain nomenclature:*

Guinot et al.^3^ argue that a major source of error in our dataset stems from how occurrences with open or uncertain nomenclature (taxon names containing qualifiers such as *cf.*, *aff.*, or *?*) are treated. In our filtered dataset, uncertain nomenclature represents 7% of all occurrences, with only 60 taxa (3%) represented exclusively by occurrences with uncertain nomenclature. In their view, these occurrences artificially extend the stratigraphic ranges of some taxa and may create spurious survivorship across the K/Pg boundary. We acknowledge that occurrences reported under open nomenclature signal taxonomic uncertainty and do not constitute definitive species-level identifications. Such uncertainty is important, particularly when these records affect temporal or geographic ranges. However, open nomenclature does not necessarily imply that the material is identifiable only at genus level, nor that it represents a different taxon. In palaeontological practice, qualifiers such as “cf.” and “aff.” can reflect preservational limitations, incomplete material, or conservative identification standards, even when the material is considered closely comparable to the nominal species. For the purposes of data integration across heterogeneous sources, we retained these occurrences under their nominal designations in the original dataset, applying this treatment uniformly. The sensitivity of our results to this decision can be directly assessed by analyses excluding such records.

*Inclusion of occurrences “without evidence”:*

Guinot et al.^3^ argue that a great number of occurrences in our dataset lack “evidence” and contend they should be excluded from analyses. Indeed, these records represent a lower-confidence class of occurrence data and their impact should be evaluated explicitly. The “evidence” terminology originates from the Paleobiology Database framework, whereby references classified as “with evidence” are those providing formal descriptions, figures, or phylogenetic justification, whereas works that list taxa without such documentation are labelled “without evidence”^4^. We consider the latter to remain usable sources of occurrence information for some large-scale analyses (indeed, they are routinely included in large-scale paleobiology studies; e.g., ^5-7^) provided their influence is documented and tested.

Both fossil and modern occurrence datasets are designed to aggregate published evidence of taxon presence through space and time, rather than to reproduce taxonomic descriptions. In palaeontology, many occurrences are reported in faunal lists or assemblage summaries (e.g., ^8-11^) and excluding such records would not only remove a substantial portion of the published fossil record, but would disregard a whole body of peer-reviewed literature. In FINS v2 these occurrences represent ~40% of the data. Excluding over 11,000 occurrences would remove a substantial portion of the available record, so it is important to evaluate whether these records alter the inferred macroevolutionary patterns rather than assume their effect *a priori*. For instance, modern biodiversity datasets routinely rely on occurrence records that lack formal taxonomic documentation, yet remain essential for ecological and macroevolutionary analyses (e.g., ^12^). If one were to require every modern occurrence to be backed by a formal description and illustration in the source publication, then a very large share of biodiversity science would collapse, because many valid occurrence records are based on monitoring schemes, survey data, or vetted human observations rather than taxonomic monographs (e.g., the Global Biodiversity Information Facility, GBIF).

*Taxonomic verification difficult due to absence of author names:*

Guinot et al.^3^ raise concerns that the absence of taxonomic authorities associated with species names may hinder taxonomic verification. We acknowledge that author names and dates can facilitate verification, particularly where the same binomen has been used for different nominal species. However, in the FINS dataset, each occurrence is directly linked to its primary literature source, allowing every record to be traced back to the publication in which the identification was made, thereby ensuring transparency and verifiability^1^. Importantly, taxonomic authorities are accessible in the Shark References website (<https://shark-references.com>). Consequently, while author names can serve as an additional layer of information, their absence in the occurrence table does not prevent taxonomic traceability because both the original literature source and synonymy information are available.

*Incorrect age assigned to some collections:*

Guinot et al.^3^ highlight a number of occurrences with erroneous age assignments. From their list, we found that seven collections and one occurrence were indeed erroneous (Table S1). While we tried to be diligent when extracting the data from publications, we were unable to completely avoid human error. As such, we have corrected these entries, resulting in the removal of 202 occurrences. Nevertheless, although Guinot et al.^3^ list occurrence PBDB_12328 as invalid based on age assignment, our assessment revealed this opinion was unfounded, with source publication providing appropriate and sufficient evidence to support the current age assignment (Table S1). As such, this occurrence was left as originally presented.

*Essential species missing resulting from excluding pre-1970 literature:*

Guinot et al.^3^ list 14 species missing from K/Pg victims and survivors. They deemed this putative gap to the focus of the FINS dataset on works published after 1970 when extracting data from the literature. Indeed, FINS combines data extracted from the literature published from 1970 onwards with data downloaded from PBDB, to which no time cut-off was imposed. The rationale behind this decision was to optimise data extraction given that the last 50 years cover the vast majority of neoselachian reports^1^. After checking these putative absences, we found that this statement was exaggerated. Indeed, eight out of these 14 species are in fact, included in FINS (Table S1), but were not included in the analyses due to the age ranges of their occurrences being too broad^2^, or were included under different valid names. Critically, only two out of the 14 “missing” species come from papers before 1970. We have added the missing four species to the dataset (Table S1).

*Taxonomic errors:*

Guinot et al.^3^ list two taxonomic misspellings which were corrected (Table S1), as well as three genera that were assigned to incorrect orders, two of which were corrected. However, we found no evidence suggesting *Squatirhina* should be placed in Rhinopristiformes. Accordingly, we have left it as originally proposed (Table S1). Guinot et al.^3^ also highlight three duplicated taxa due to incorrect taxonomy, which we have also harmonised throughout (Table S1), as well as four disused genera that were standardized following current taxonomic treatments. Finally, Guinot et al.^3^ flag additional taxonomic issues for 15 taxa. In five of these cases, the issues were due to updates in Shark References and PBDB that were not captured at the time of the data collection. In a small number of cases, records were marked invalid or removed due to stratigraphic inconsistencies, uncertain identification, or confirmed database errors. Other flagged cases were reviewed but required no change because they represent either valid taxonomic treatments in current references or differences in taxonomic interpretation.

**Supplementary methods**

***DeepDive analyses***

We performed DeepDive analyses of the two new datasets (at the species level with the two data treatments described in the main text and Table S2). We followed the same analysis as Gardiner et al^2^. Accordingly, we generated 100 replicates of the datasets while resampling the ages of the occurrences from their stratigraphic ranges. Using DeepDive’s simulation autotuning function we generated for each dataset 60,000 datasets, 80% of which were used for training while the remaining 20% served as validation set. We trained four models with different architectures from which we obtained an ensemble of predicted diversity trajectories as in^2^.

To further test whether the models might overlook a mass extinction at the K/Pg we ran an additional set of analyses explicitly trained for this purpose. In this case we customized the DeepDive simulation setting to enforce, in all training simulations, a fixed extinction event at 66 Ma with magnitude drawn from a uniform distribution from 0 to 99%. This ensures that a substantial fraction of the training datasets (~ one in five) were generated with an extinction event removing more than 80% of standing diversity at the K/Pg boundary. We used these trained models to predict neoselachian diversity and found that the predictions did not substantially shift under these models (Figure 1).

**Supplementary Figures**

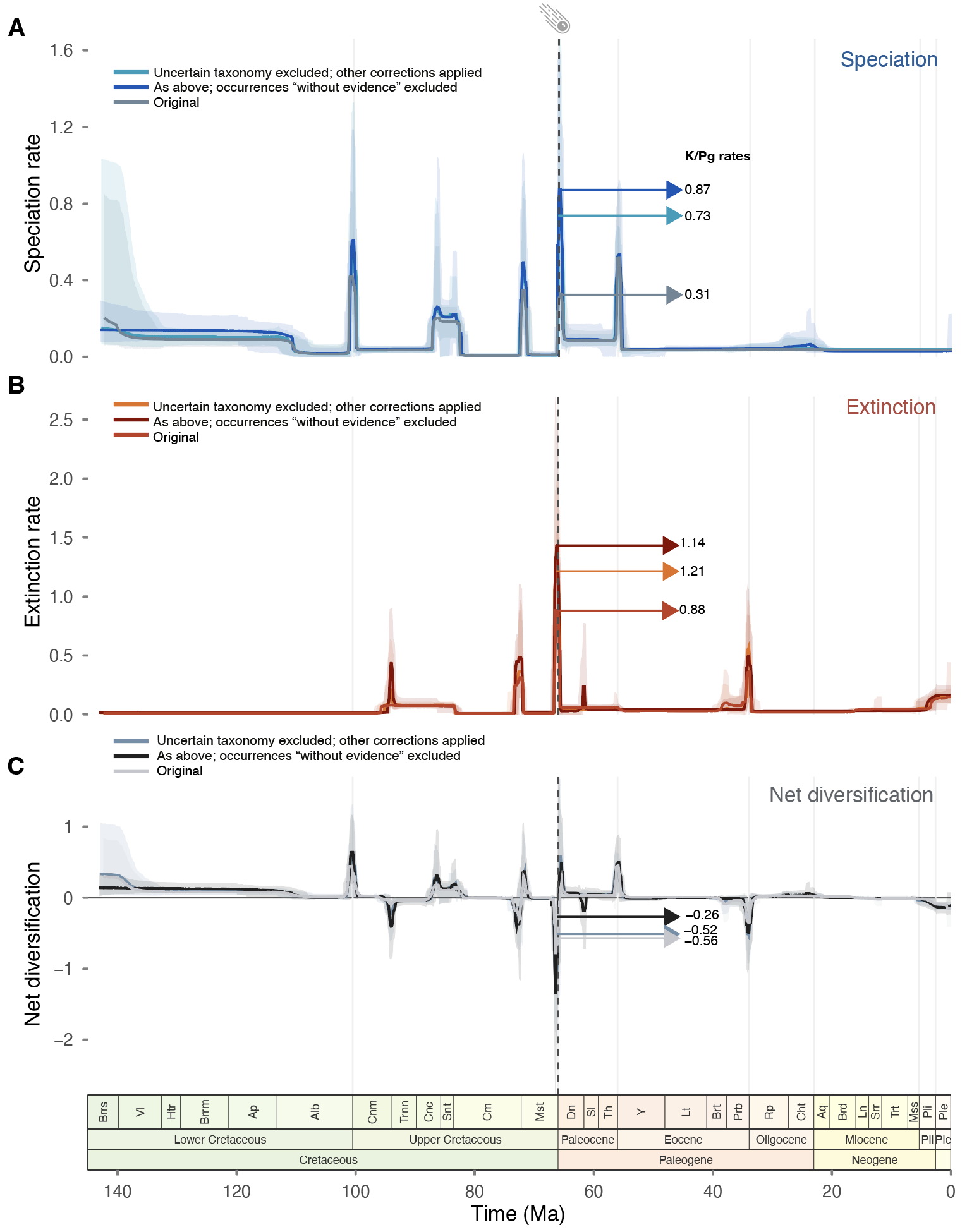

**Figure S1. Diversification trajectories of neoselachian species global assemblage from the Early Cretaceous to the Pleistocene**. **A.** Speciation, **B.** Extinction, **C.** Net diversification rates. Solid lines represent the mean posterior rates and shaded areas the 95% CIs. Net diversification is the difference between speciation and extinction rates. Abbreviations: Pli. = Pliocene, Ple. = Pleistocene, Ber. = Berriasian, Val. = Valanginian, Hau. = Hauterivian, Bar. = Barremian, Apt. = Aptian, Alb. = Albian, Cen. = Cenomanian, Tur. = Turonian, Con. = Coniacian, San. = Santonian, Cam. = Campanian, Maa. = Maastrichtian, Dan. = Danian, Sel. = Selandian, Tha. = Thanetian, Ypr. = Ypresian, Lut. = Lutetian, Bart. = Bartonian, Pri. = Priabonian, Rup. = Rupelian, Cha. = Chattian, Aqu. = Aquitanian, Bur. = Burdigalian, Lan. = Langhian, Ser. = Serravallian, Tor. = Tortonian, Mes. = Messinian. Dashed vertical line and meteorite shape denote the Cretaceous/Paleogene (K/Pg) extinction event. Horizontal arrows indicate rates at the K/Pg.

**Supplementary Tables**

**Table S1. Audit of “corrections” proposed by Guinot et al.^3^** An assessment of each data issue reported in both their main manuscript and its supplementary material.

| **Type of issue flagged** | **Issues flagged** | **Response** | **Action** |
| --- | --- | --- | --- |
| Incorrect age | Collections L_1416–L_1419 | Reworked material^13^. Age marked as invalid for 4 collections and 132 associated occurrences. | Corrected |
|  | Collection L_3034 | Identified as reworked material^14^. Age marked as invalid for the collection and 23 associated occurrences. | Corrected |
|  | Occurrence PBDB_17257 *Onchopristis numidus* | Age assignment was deemed uncertain^15^, and the occurrence age was marked as invalid. | Corrected |
|  | Occurrence PBDB_12328 – *Myliobatis wurnoensis* | Source publication does not indicate age uncertainty, instead, reports this as the first Cretaceous occurrence of the species without questioning its stratigraphic context^15^. | None |
|  | Collection PBDB_214834 | Guinot et al.^3^ suggest that this represents reworked material. Pending verification of the source publication^16^, the collection age and its 23 associated occurrences were provisionally marked as invalid. | Corrected |
|  | Collection PBDB_214373 | Same as PBDB_214834. | Corrected |
| *Essential species missing* | *Chiloscyllium broennimanni* (Casier, 1958)^17^ | Occurrences from Casier (1958)^17^ are included in FINS from PBDB, but this species is not present in this source. The species is present in FINS based on Kruckow & Thies (1990)^18^, but the occurrence has an age range too broad (36 Myr) to meet the analytical filtering criteria (occ. L_5949). | None |
|  | *Cretolamna maroccana* (Arambourg, 1935)^19^ | Treated as a synonym of *Serratolamna maroccana* in FINS. There are 21 occurrences in FINS, 18 of which were also present in the filtered dataset originally analysed (Data S1 in Gardiner et al., 2026^2^). | None |
|  | *Erguitaia arganiae* (Arambourg, 1952)^20^ | This species was absent from FINS v1 and an occurrence (occ. L_17915) had been already added from Cappetta (1989)^21^ in FINS v2, which is published along with this study and included in sensitivity tests . | None |
|  | *Angolabatis benguelaensis* (Antunes & Cappetta, 2002)^22^ | Present in FINS (occ. L_5007), but excluded from analyses due to broad age range (17.6 Myr). | None |
|  | *Australopristis wiffeni* (Martill & Ibrahim, 2012)^23^ | Rather than reporting new occurrences, Martill & Ibrahim (2012)^23^ reinterpret specimens previously reported as *Onchopristis numidus^24^*. Occurrences of this species already exist in FINS (PBDB_23418, PBDB_23950). A “reidentification” column was added and the taxonomy was updated. However, they will be excluded from the filtered dataset due to broad age ranges (17.6 Myr). | Adjusted |
|  | *Carcharias heathi* (Case & Cappetta, 1997)^25^ | Present in FINS (occ. PBDB_25868) but considered a synonym of *Carcharias samhammeri.* There are 77 occurrences in FINS and 65 were present in the filtered dataset. | None |
|  | *Chlamydoselachus gracilis* (Antunes & Cappetta, 2002)^22^ | Present in FINS (occ. L_4995) but excluded due to broad age range (17.6 Myr). | None |
|  | *Cretascymnus quimbalaensis* (Antunes & Cappetta, 2002)^22^ | Present in FINS (occ. L_50) but excluded due to broad age range (17.6 Myr). | None |
|  | *Hamrabatis ornata* (Cappetta, 1991)^26^ | Real omission. Publication is now incorporated into FINS (as Cappetta 1991b)^26^ - three collections (PBDB_275564 to PBDB_275566) and 58 occurrences added (PBDB_2039328 to PBDB_2039385). | Corrected |
|  | *Hispidaspis turkestanensis* (Zhelezko, 2000)^27^ | Real omission. Publication is now incorporated into FINS (as Zhelezko 2000b)^27^ - seven collections (L_3471 to L_3477) and 14 occurrences added (L_30505 to L_30518) | Corrected |
|  | *Ischyrhiza monasterica* (Case & Cappetta, 1997)^25^ | Present in FINS and the filtered dataset as a synonym of *Ischyrhiza avonicola* (occ. PBDB_25884). | None |
|  | *Ptychotrygon greybullensis* (Case, 1987)^28^ | Present in FINS and the filtered dataset as synonym of *Texatrygon hooveri* (4 occurrences – PBDB_26648, PBDB_26747, PBDB_19140, PBDB_19213) | None |
|  | *Scyliorhinus cepaeformis* (Halter, 1990)^29^ | Real omission. Halter (1990)^29^ is now incorporated into FINS - 42 occurrences added (L_30448 to L_30489) | Corrected |
|  | *Scyliorhinus wardi* (Halter, 1990)^29^ | Same as above. | Corrected |
| Taxonomic issues (misspellings) | *Dasyatis* sp. | Occ. L_29378 was mistakenly entered as cf. *Dasyatis* sp. Rafinesque into FINS. It was changed to *Dasyatis* sp. and the rank was changed to genus. | Corrected |
|  | *Pararhincodon cf. crochardi* | Occ. L_21547 was originally entered as *Pararhincodon crochardi*, now corrected to *Pararhincodon cf. crochardi.* | Corrected |
| Taxonomic issues (order assignments) | *Potamotrygon* should be Myliobatiformes | The assignment of *Potamotrygon* to Rajiformes reflects an earlier version of FINS and has since been corrected to Myliobatiformes; this affects 23 occurrences. | Corrected |
|  | *Squatirhina* should be in Rhinopristiformes | The assignment to Rajiformes follows the classification adopted in Shark-References and is consistent with published works (e.g., ^30,31,32^). | None |
|  | *Belemnobatis* should be in Apolithabatiformes | Some occurrences of *Belemnobatis* were incorrectly placed in Rajiformes in the version of FINS that was used in the original analyses^2^. We have updated the order classification of all occurrences of this genus to Apolithabatiformes, following Tütcher et al. (2025)^33^. We have also updated all occurrences of *Spathobatis*, which has also been assigned to Apolithabatiformes. This change is reflected in the new versions of FINS (v2). | Adjusted |
| Taxonomic issues (duplicated genera due to incorrect taxonomy) | *Cretolamna/Cretalamna* | Both are considered as valid names in the Shark References backbone, each associated with a unique species. Although Cappetta (2012)^34^ states that only *Cretolamna* should be used, *Cretalamna* has also been used after 2012 (e.g., ^35,36^). Nevertheless, *Cretalamna* was added as synonym of *Cretolamna* in our synonym table and all *Cretalamna* instances were accordingly updated to *Cretolamna*, affecting 54 occurrences. | Corrected |
|  | *Notidanus/Hexanchus* | *Hexanchus* was added as a synonym to *Notidanus* and all instances of *Notidanus* were updated accordingly (11 occurrences). | Corrected |
|  | *Rhinobatos/Rhinobatus* | *Rhinobatus* was added as a synonym to *Rhinobato*s and all instances of *Rhinobatus* were updated accordingly (7 occurrences).  We also updated the accepted name of *Rhinobatus expansus* (occ. PBDB_22901) to *Pararaja expansus* following Cappetta (2012)^34^ and updated the synonym table. | Corrected |
| Taxonomic issues (disused taxa) | *Eorhincodon* | *Eorhincodon tianshanensis* was updated to *Rhizoprionodon tianshanensis* following Cappetta (2012)^34^, and the genus *Eorhincodon* was otherwise treated as a synonym of *Pseudomegachasma* based on Shimada et al. (2015)^37^. During this revision, two occurrences originally entered as *Eorhincodon* sp. were corrected to *Eorhincodon casei* with the valid name *Pseudomegachasma casei* (although these records have invalid ages and are therefore not present in the filtered dataset analysed). | Corrected |
|  | *Hypolophus, Oxyrhina* and *Orthacodus* | Additional genus-level harmonisations included treating *Hypolophus* as a synonym of *Hypolophodon* (3 occurrences), *Oxyrhina* as a synonym of *Isurus* (10 occurrences), with the species *Oxyrhina acuminata* further resolved to *Paranomotodon angustidens,* and *Orthacodus* as a synonym of *Sphenodus* (5 occurrences). | Corrected |
| Taxonomic issues (specific cases) | *Paratriakis curtirostris* | Previously considered valid, but now considered a synonym of *Palaeotriakis curtirostris* (10 occurrences affected). | Corrected |
|  | *Lamna mediavia* | Previously considered valid, but now considered a synonym of *Otodus obliquus*. Family classification was corrected from Lamnidae to Otodontidae. | Corrected |
|  | *Prosopodon* | Although this genus remains valid in the current Shark References taxonomic backbone, Cappetta (2012)^34^ suggested replacing it with *Phosphatodon* (6 occurrences affected) | Corrected |
|  | *Striatolamia apiculatus* | Guinot et al.^3^ proposed this to be a synonym of *Cosmopolitodus hastalis*, however based on the Shark References synonym dictionary it is a valid name. The issue with this occurrence concerns stratigraphic age rather than taxonomy. The specific occurrence flagged by Guinot et al.^3^ (occ. PBDB_12099) was far outside of the age range from all other occurrences of this species and was therefore marked as invalid (1 occurrence affected). | Corrected |
|  | *Myliobatis dixoni* | The occurrence PBDB_12894 was flagged due to it being the only occurrence of *M. dixoni* from the Cretaceous. After evaluation, this occurrence was deemed invalid as its age was far outside of the age range from all other occurrences of this species. This outlier occurrence was reported as *Myliobatis obesus* in its source publication, which was considered to be a synonym of *M. dixoni,* however both are now considered synonyms of *Aetomylaeus dixoni*. The synonym table and all relevant occurrences were updated accordingly (96 occurrences updated in the taxonomy and 1 occurrence invalidated). | Corrected |
|  | *Myliobatis* (Cretaceous occurrences) | Guinot et al.^3^ stated that the genus should be restricted to the Cenozoic. Most occurrences are identified only to genus level, three occurrences are identified to the species level – *Myliobatis leidyi* was entered as *Myliobatis cf. leidyi* and will be excluded from the sensitivity tests, however we did not find a reason to exclude the other two -*Myliobatis wurnoensis* is discussed above, *Myliobatis foxhillsensis* is considered a valid species and its age is not disputed by the authors (Hoganson et al. 2019)^38^; 10 occurrences reviewed). | None |
|  | *Lamna* (Cretaceous occurrences) | Guinot et al.^3^ state that the genus *Lamna* should be restricted to extant and Cenozoic taxa without providing any evidence. One species-level occurrence (*Lamna marginalis*) from the Cretaceous is valid according to Shark References and the remaining occurrences are genus-level (13 occurrences reviewed). | None |
|  | *Notidanodon* | Taxonomic revision indicates that some Cenozoic species previously placed in *Notidanodon* belong in the genus *Xampylodon*. The synonym table and relevant occurrences were updated so that *Notidanodon brotzeni* and *Notidanodon loozi* are treated as *Xampylodon brotzeni* and *Xampylodon loozi* (19 occurrences updated). | Corrected |
|  | *Notidanodon* (genus-level occurrences) | Two occurrences reported as *Notidanodon* sp. were reviewed and considered taxonomically invalid based on the discussion in the source publication^39^, where the material was described as damaged and intermediate between species (occ. PBDB_10310 and PBDB_10319). The occurrences were therefore marked invalid. Three occurrences of *Notidanodon* sp. from the Cenozoic remained marked as valid after our revision (5 occurrences reviewed). | Corrected |
|  | *Serratolamna aschersoni* | Guinot et al.^3^ highlighted that occurrences of *Cretolamna aschersoni* were given the valid name of *Serratolamna aschersoni*, expanding the range of the typically Cretaceous genus *Serratolamna* into the Cenozoic. Indeed, in the previous version of FINS *C. aschersoni* was treated as a synonym of *S. aschersoni,* however the current version of Shark References shows the opposite relationship (i.e., *S. aschersoni* is a synonym of *C. aschersoni*). We updated the synonym table and occurrences accordingly (12 occurrences affected). | Corrected |
|  | *Somniosus* / *Rhinoscymnus* | An issue in the synonym table caused species-level occurrences originally identified as *Rhinoscymnus* to have their genus automatically reassigned to *Somniosus*. This was corrected so that species-level occurrences retain the genus *Rhinoscymnus* where appropriate (3 occurrences corrected). | Corrected |
|  | *Heptranchias* | An occurrence included in the previous version of FINS from PBDB was later removed from PBDB because it had been entered under an incorrect name in the UCMP collection database (https://ucmpdb.berkeley.edu/). The occurrence was therefore removed from FINS (1 occurrence affected). | Corrected |
|  | *Palaeohypotodus* / *Odontaspis bronni* | Guinot et al.^3^ suggested that treating the Maastrichtian occurrence of *Odontaspis bronni* (occ. PBDB_16301) as a synonym of *Palaeohypotodus bronni* is not typical. However, the current Shark References taxonomy supports the synonymy used in FINS and recent publications (e.g.,^40^) also recognize this specific occurrence from Cappetta and Corral (1999)^41^ as a synonym of *P. bronni*.(1 occurrence reviewed). Furthermore, Guinot et al.^3^ highlighted the genus *Palaeohypotodus* as a typically Cenozoic genus with problematic Cretaceous occurrences, however this genus is found in both eras (e.g., ^40^). | None |
|  | *Synechodus subulatus* | The Paleogene occurrence of this species (occ. L_21736) was erroneously entered into FINS as *Synechodus subulatus,* whereas in the source publication^42^ it was presented as *Synechodus aff. subulatus*. This entry error was fixed, resulting in exclusion of this occurrence from the sensitivity analyses (1 occurrence affected). | Corrected |
|  | *Mesiteia emiliae* | We found no indication in the original publication Carnevale et al. (2014)^9^ questioning the identification of the occurrence (occ. PBDB_10599). | None |

**Table S2. Data treatment and methodological approach for sensitivity tests.** Two data subsets were created and three sensitivity analyses were performed to assess the impact of Guinot et al.’s “corrections”^3^ in our results.

| **Approach** | | | **Original** | **Test 1** | **Test 2** | **Test 3** |
| --- | --- | --- | --- | --- | --- | --- |
| Occ with uncertain nomenclature | | | Included | Excluded | Excluded | Excluded |
| Occ “without evidence” | | | Included | Included | Included | Excluded |
| DeepDive K/Pg model | | | Standard | Standard | Explicitly modelled | Standard |
| Species level | Total occ | | 18,328 | 16,767 | 16,767 | 10,812 |
|  | Range through (raw) mean diversity | Diversity Maastrichtian/Danian | 221/150 | 207/119 | 207/119 | 174/90 |
|  |  | Diversity change | -32% ± 2% | -42% **±** 2% | -42% **±** 2% | -48% **±** 3% |
|  | DeepDive mean diversity | Diversity Maastrichtian/Danian | 1420/1283 | 1404/1083 | 1564 /1099 | 1744 /1166 |
|  |  | Diversity change | **-10%±9%** | **-22.9±1.1%** | **-29.7±8.4%** | **-33.1±3.9%** |
